# Identification of bacteriophage genome sequences with representation learning

**DOI:** 10.1101/2021.09.25.461359

**Authors:** Zeheng Bai, Yao-zhong Zhang, Satoru Miyano, Rui Yamaguchi, Kosuke Fujimoto, Satoshi Uematsu, Seiya Imoto

## Abstract

**Motivation:** Bacteriophages/Phages are the viruses that infect and replicate within bacteria and archaea, and rich in human body. To investigate the relationship between phages and microbial communities, the identification of phages from metagenome sequences is the first step. Currently, there are two main methods for identifying phages: database-based (alignment-based) methods and alignment-free methods. Database-based methods typically use a large number of sequences as references; alignment-free methods usually learn the features of the sequences with machine learning and deep learning models.

**Results:** We propose INHERIT which uses a deep representation learning model to integrate both database-based and alignment-free methods, combining the strengths of both. Pre-training is used as an alternative way of acquiring knowledge representations from existing databases, while the BERT-style deep learning framework retains the advantage of alignment-free methods. We compare INHERIT with four existing methods on a third-party benchmark dataset. Our experiments show that INHERIT achieves a better performance with the F1-score of 0.9932. In addition, we find that pre-training two species separately helps the non-alignment deep learning model make more accurate predictions.

**Availability:** The codes of INHERIT are now available in: https://github.com/Celestial-Bai/INHERIT.

**Supplementary information:** Supplementary data are available at *BioRxiv* online.

## 1 Introduction

Bacteriophages (phages for short) are the viruses that infect bacteria and archaea, and are rich in human body (Fuhrman, 1999; Edwards and Rohwer, 2005; Rohwer and Thurber, 2009; Rodriguez-Valera *et al*., 2009; Reyes *et al*., 2012). To study the role of phages in microbial community in the human body, we need first to identify phages from the metagenome nucleotide sequences (Marquet *et al*., 2020; Fang *et al*., 2019). Using a method that can precisely distinguish between phages and bacteria can help researchers study phages more efficiently. Many methods have been proposed to identify phages, such as VIBRANT (Kieft *et al*., 2020), VirSorter2 (Guo *et al*., 2021), Seeker (Auslander *et al*., 2020) and DeepVirFinder (Ren *et al*., 2020). We categorize them into two groups: database-based (alignment-based) methods (VIBRANT and VirSorter2), and alignment-free methods (Seeker and DeepVirFinder). Both types have their advantages and disadvantages, and they are complementary. Database-based approaches are commonly based on multiple sequence alignment (Edgar and Batzoglou, 2006; Hyatt *et al*., 2010; Chatzou *et al*., 2016) with Profile Hidden Markov Models (Eddy, 1998), which can achieve good prediction performance. However, such prediction speed is generally limited by alignment. Alignment-free methods usually can predict fast. However, subject to the training process of the machine learning and deep learning models, we need to balance the amount of phage and bacteria data (Japkowicz and Stephen, 2002), which affects the amount of information obtained. The introduction of them can be found in Section 1.1, Supplementary Methods.

In proposing the MSA Transformer, Rao *et al*. (2021) demonstrated that pre-trained Transformer-based models can have comparable performance to HMM Profiles and are even better in some cases. That indicates the core of the database-based approaches, HMM Profiles, can be realized for a similar purpose by representation learning. Thus we can use the pre-train-fine-tune paradigm (Liu *et al*., 2021; Mao, 2020; Gururangan *et al*., 2020; Radford *et al*., 2018; Bengio *et al*., 2013; Zhang *et al*., 2018) to combine the above two methods.

Here we propose INHERIT: **I**dentificatio**N** of bacteriop**H**ag**E**s using deep **R**epresentat**I**on model with pre-**T**raining. It also means our model “inherits” the characteristics from both database-based and alignment-free methods. The code of INHERIT is now available at https://github.com/Celestial-Bai/INHERIT. We show that using the representation learning framework can improve deep learning models, and INHERIT also achieves the best performance in our tests.

The main contributions of our paper can be summarized as follows:

1. We proposed INHERIT, an integrated model based on the DNA sequence language model: DNABERT, with two pre-trained models as references. It reaches the best performance compared with four existing state-of-the-art approaches: VIBRANT, VirSorter2, Seeker, and DeepVirFinder. INHERIT outperforms them with the highest F1-score of 0.9932 in our test.
2. We used an independent pre-training strategy to deal with the data imbalance issue of bacteria and phages. We found that this strategy can help the deep learning framework make more accurate predictions for both species.

## 2 Methods

INHERIT is a model with DNABERT as the backbone and uses two pre-trained models as references (see the pipeline in Figure 1A). DNABERT is an extension of BERT (Devlin *et al*., 2018) on nucleotide sequences. The structure of BERT model contains of 12 Transformer encoders (Vaswani *et al*., 2017), and Transformer is a neural network composed mainly of multi-head self-attention. Multi-head self-attention is a mechanism that can be expressed with (Vaswani *et al*., 2017):

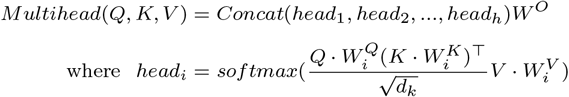

**Fig. 1:**
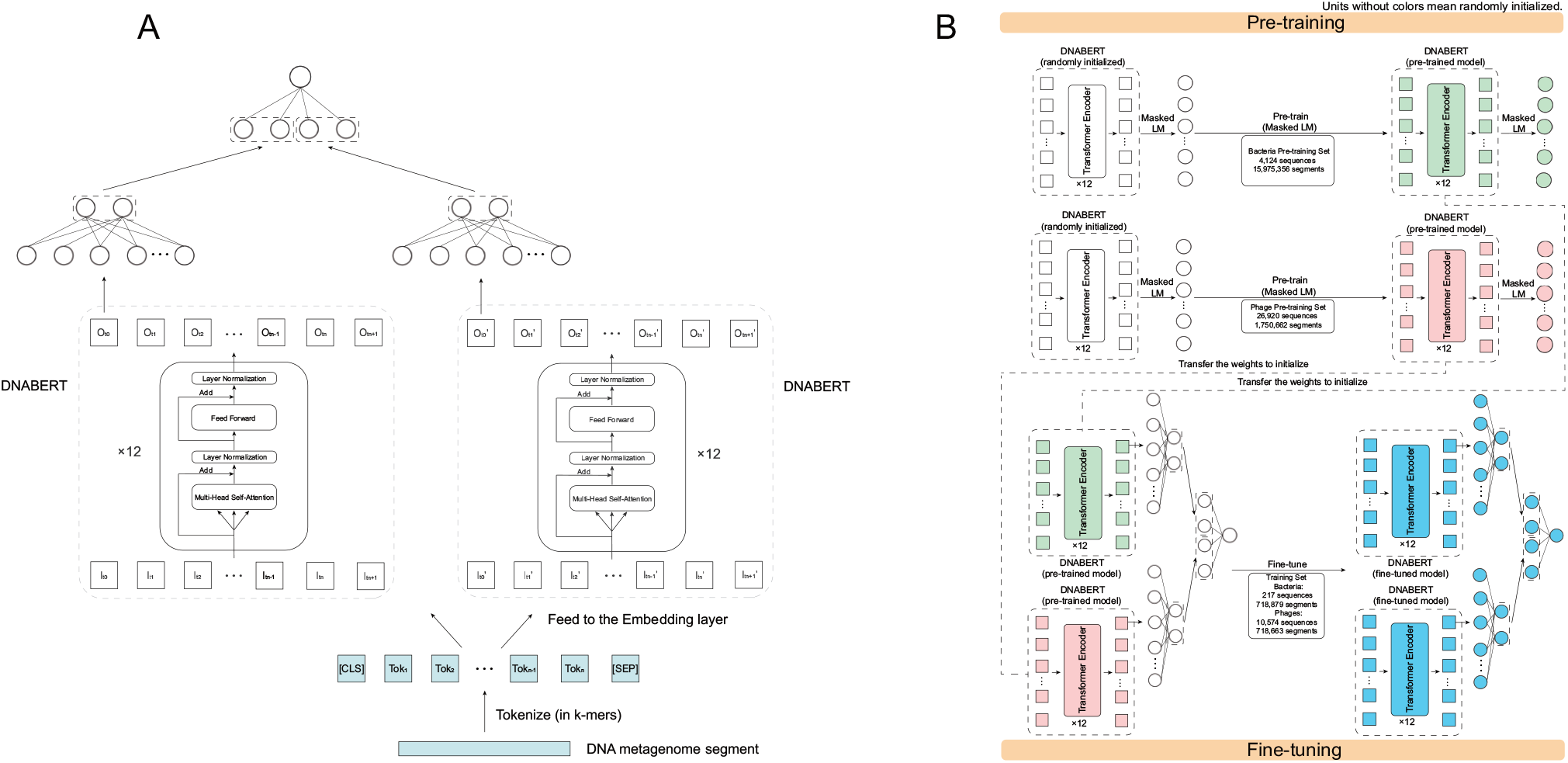
(A) shows the overall model architecture and explains how the label class is predicted given trained model parameters. In predicting the label class of a given 500 bp nucleotide fragment, it is first encoded into tokens as the k-mer inputs. Then, they will be converted to the embeddings for two fine-tuned DNABERTs through the embedding layer independently. These embeddings are then fed into the fine-tuned Transformer encoders. The representations of the [CLS] of the two DNABERTs are extracted and then classified to generate four outputs. Eventually, INHERIT predicts the class from the four outputs of DNABERTs with a dense layer. (B) shows how the parameter weights are assigned through pre-training and fine-tuning. In pre-training, two randomly initialized DNABERTs pre-train independently with two datasets of different sizes and species. After that, the pre-trained weights in the lower non-linear-probing layers are first used for the model initialization. Then in the fine-tuning step, all model parameters, including those of the two pre-trained DNABERTs, are fine-tuned together with a balanced training set.

Where *Q, K, V* are the vectors obtained by multiplying the three learned vectors with the last hidden states. 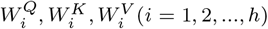 and *W*^*O*^ are all learned matrices. *h* is the number of attention heads, and *d*_*k*_ is the dimension of *K*. A detailed introduction of DNABERT can be found in the Supplementary Methods Section S1.1. We fine-tune the two pre-trained models simultaneously to identify the metagenome sequences. The following will introduce the structure of INHERIT and the datasets we use.

### 2.1 Model architecture and pipeline

Here we show the architecture of INHERIT in Figure 1A and how parameter weights are assigned and transferred in the pre-training and fine-tuning process.

The sequences are split into several 500 bp-long segments as the input of INHERIT. When this sequence is not divisible by 500, we will use the head of this sequence to complement its end until it is divisible, which keeps the same with Seeker. For each segment, it will be encoded to tokens as k-mer inputs. According to the previous work (Ji *et al*., 2021), we use 6-mer as input to DNABERT. Those tokens will be generated to embeddings for two fine-tuned DNABERTs through each embedding layer separately. Then each embedding is fed to each transformer encoder (Vaswani *et al*., 2017). Same with BERT (Devlin *et al*., 2018), the representations of the “[CLS]” token is extracted and generate two outputs for each fine-tuned DNABERT through a dense layer (Wolf *et al*., 2020). INHERIT eventually predicts the label class based on the four outputs with a dense layer. The prediction of the whole sequence is the average of predictions of all segments, which we call the “score” of the sequence.

Here we used the pre-train-fine-tune paradigm to build INHERIT (see the process of pre-training and fine-tuning INHERIT in Figure 1B). To deal with the information bias that may be caused by data imbalance (Thabtah *et al*., 2020), we pre-trained bacteria and phages independently. The number of bacteria we have known is much larger than the number of phages, and the length of bacteria is also longer (Chanishvili *et al*., 2001). If we want the pre-training set to carry a considerable amount of data, the segments belonging to the bacteria will be bound to be much more than those belonging to the phages. If we combine bacteria and phages in one pre-trained model, the model will learn much more about bacteria than phages. Therefore, we prepared two pre-trained models for INHERIT. Here, we used Masked Language Modeling (Devlin *et al*., 2018; Naseem *et al*., 2021) as the pre-training task, which is the same as Ji *et al*. (2021). For the detailed settings of the pre-training, please refer to Supplementary Methods Section S1.2.

Fine-tuning is very similar to the traditional training strategy (Dodge *et al*., 2020). The difference between them is that traditionally we randomly initialize the deep learning model before we start to train the model. However, we will transfer most of the pre-trained model weights as the initialization before we start to fine-tune the deep learning framework. Here we transferred the weights of non-linear-probing layers of the two pre-trained models to initialize INHERIT. The linear layers of INHERIT were still randomly initialized because we could not transfer the weights from the pre-trained models. All model parameters were fine-tuned together with a balanced training set. For hyperparameters and platforms of fine-tuning, please see Supplementary Methods Section S1.2.

### 2.2 Datasets

#### 2.2.1 Pre-training sets

To make pre-trained models carry as much biological information as possible, we pre-trained bacteria and phages separately and did not balance the size of the two pre-training sets. For the bacteria pre-training set, we used ncbi-genome-download (https://github.com/kblin/ncbi-genome-download) to obtain the complete bacteria genome sequence from the NCBI FTP. We used the command: ncbi-genome-download-formats fasta –assembly-levels complete bacteria. All of those bacteria sequences were high quality, and we called them “bacteria assemblies”. We randomly sampled 4,124 bacteria sequences from them because of the physical memory limitation. However, these 4,124 sequences can generate 15,975,346 segments, and the dataset size is large enough. We could not obtain the phage sequence data in the same way for the phage pre-training set. Since phage sequences could not be found and downloaded directly in the NCBI FTP like the bacteria sequences, we directly searched for the keyword “phage” on NCBI, downloaded all sequences longer than 500 bp, and checked all of them manually. We also referred to the phage sequences used by Seeker and VIBRANT and finally generated a pre-training set containing 26,920 phage sequences. It did not include the phage sequences in the test and validation sets to prevent overfitting. These phage sequences can generate 1,750,662 segments, and the size is still large for a phage dataset.

#### 2.2.2 Training set and validation set for fine-tuning

For bacteria during fine-tuning, we randomly selected 260 bacteria sequences that were not in the pre-training and test sets but bacteria assemblies. 217 bacteria sequences were used as the training set, generating 718,879 segments, and the remaining 43 were used as the validation set, generating 188,149 segments. However, we did not have as many sequences to choose from for phages, so we selected 10,574 phage sequences from the pre-training set that possessed a quality comparable to the bacteria assemblies, generating 718,663 segments. We also chose 2,643 sequences not in the pre-training set as the validation set, generating 186,121 segments.

#### 2.2.3 Test set for comparisons

The test set we use is one of the third-party benchmark tests previously proposed by Ho et al. Ho *et al*. (2021) for virus identification methods, called the RefSeq test set. Since our method identifies phages and not other viruses, we only used data related to phages. The RefSeq test set contains 710 bacteria sequences and 1,028 phage sequences. However, since there are 19 bacteria sequences removed from NCBI RefSeq database, we used the rest of them, including 691 bacteria sequences and 1,028 phage sequences, to examine the performance of the methods on phage identification. It should be added that, in that article (Ho *et al*., 2021), the authors split the sequences in this test set into 1kb to 15kb segments on average and predict the results and calculate metrics on segment level to make a benchmark test. However, since we consider INHERIT to determine whether a sequence is a phage or not in applications, we compared INHERIT with other existing methods on the sequence level. For experimental details and results, please refer to Section 3.1.

We have posted the accessions of the sequences used in each dataset and the sources we obtain in Supplementary Table S1.

## 3 Experiments

### 3.1 Benchmarking INHERIT with VIBRANT, VirSorter2, Seeker, DeepVirFinder

In this section, we compared INHERIT with four state-of-the-art methods: VIBRANT, VirSorter2, Seeker, and DeepVirFinder, which are the representatives of database-based and alignment-free methods. We used a third-party benchmark dataset to conduct experiments and analyses.

#### 3.1.1 Experimental setups

##### Baselines

Until our work is completed, INHERIT is the only model that integrates the features of both database-based and alignment-free approaches. Thus, we chose two representatives which have achieved state-of-the-art from each of the two methods, VIBRANT (Kieft *et al*., 2020) and VirSorter2 (Guo *et al*., 2021); and Seeker (Auslander *et al*., 2020) and DeepVirFinder (Ren *et al*., 2020), to compare it with INHERIT. To ensure the comparison is as fair as possible, we used each method’s default commands for predictions as much as possible, and we set corresponding rules for some methods to ensure that the prediction formats of each method are as consistent as possible. Detailed settings for each method can be found in Supplementary Methods Section S1.3.

##### Evaluation metrics

The evaluation metrics we chose are:

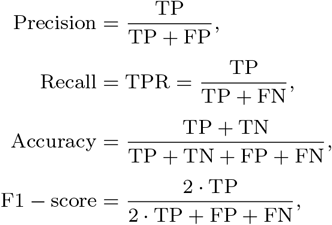

and AUROC (Area Under the Receiver Operating Characteristic curve) and AUPRC (Area Under the Precision-Recall Curve). In this paper, TP is the number of phage sequences successfully identified as phages, while FP is the number of bacteria sequences incorrectly identified as phages. TN is the number of bacteria sequences successfully identified as bacteria and FN is the number of phage sequences incorrectly identified as bacteria. AUROC and AUPRC are calculated based on the score of each sequence and the real value (phages are recorded as 1 and bacteria as 0). Here we calculate all evaluation metrics by scikit-learn v1.0.2 (Pedregosa *et al*., 2011).

##### Prediction results

The predictions of VIBRANT, VirSorter2, Seeker, DeepVirFinder, and INHERIT for all the sequences in the test set can be seen in Supplementary Table S2.

#### 3.1.2 Result and analysis

##### INHERIT performs better than existing methods

From the results (see Table 1), compared to VIBRANT, VirSorter2, Seeker, and DeepVirFinder, INHERIT performs better than other existing methods. Moreover, the overall performance of INHERIT is an order of magnitude more precise. Even if we use the default threshold of 0.5, the recall of INHERIT does not differ much from that of VIBRANT. Significantly, the high F1-score of INHERIT proves that INHERIT performs well enough when we use the default parameters and is competent for most application scenarios. From the p-values of the DeLong Test (DeLong *et al*., 1988) of every two methods, the ROC curves for each of the two approaches are statistically significantly different. Therefore, the high AUROC and AUPRC indicate that INHERIT can distinguish between phages and bacteria better.

**Table 1.** The benchmark results of VIBRANT, VirSorter2, Seeker, DeepVirFinder and INHERIT. For all methods we used the default commands for predictions. This can make our benchmarking as fair as possible. Values corresponding to best performance are bolded.

| Model | Precision | Recall | Accuracy | F1-score | AUROC | AUPRC |
| --- | --- | --- | --- | --- | --- | --- |
| VIBRANT | 0.9541 | 0.9903 | 0.9656 | 0.9718 | 0.9595 | 0.9751 |
| VirSorter2 | 0.9685 | 0.9893 | 0.9728 | 0.9787 | 0.9932 | 0.9971 |
| Seeker | 0.9264 | 0.8453 | 0.8674 | 0.8840 | 0.9382 | 0.9605 |
| DeepVirFinder | 0.8992 | 0.8502 | 0.8534 | 0.8740 | 0.8971 | 0.9435 |
| **INHERIT** | **0.9884** | **0.9981** | **0.9919** | **0.9932** | **0.9996** | **0.9997** |

##### The performance of INHERIT is less affected by the length of nucleotide sequences

We find the performance of INHERIT is less sensitive to the length of genome sequences compared with other methods. We divided the phage and bacteria into five length intervals, each based on the quartiles of lengths in the test set. The length intervals are according to the following rules: “minimum - first quartile”, “first quartile - median”, “median - third quartile”, “third quartile - maximum of non-outliers”, “the maximum of non-outliers - outliers”. For phage sequences, there are “less than 42,000 bp”, “42,000 bp - 51,000 bp”, “51,500 bp - 91,600 bp”, and “greater than 91,600 bp”. For bacteria sequences, there are “less than 2,788,000 bp”, “2,788,000 bp - 4,107,500 bp”, “4,107,500 bp - 4,938,000 bp “, “greater than 4,938,000 bp”. We calculated the true positive rates of VIBRANT, VirSorter2, Seeker, DeepVirFinder, and INHERIT in the four intervals of phage and the true negative rates in the four intervals of bacteria, respectively. That allows exploring whether these five methods perform consistently and robustly at different sequence lengths for both phages and bacteria. We can find that INHERIT is the only method to maintain the true positive rate of 0.99 and the true negative rate of 0.97 in different intervals (see Figure 2). It does not always perform the best in every interval, but it is more robust than other models. For example, the true negative rate of VIBRANT in the “2788000 bp-4107500 bp” interval is 0.994, which is better than INHERIT (0.977), but its performance in the “4938000 bp-8000000 bp” interval is only 0.767.

**Fig. 2:**
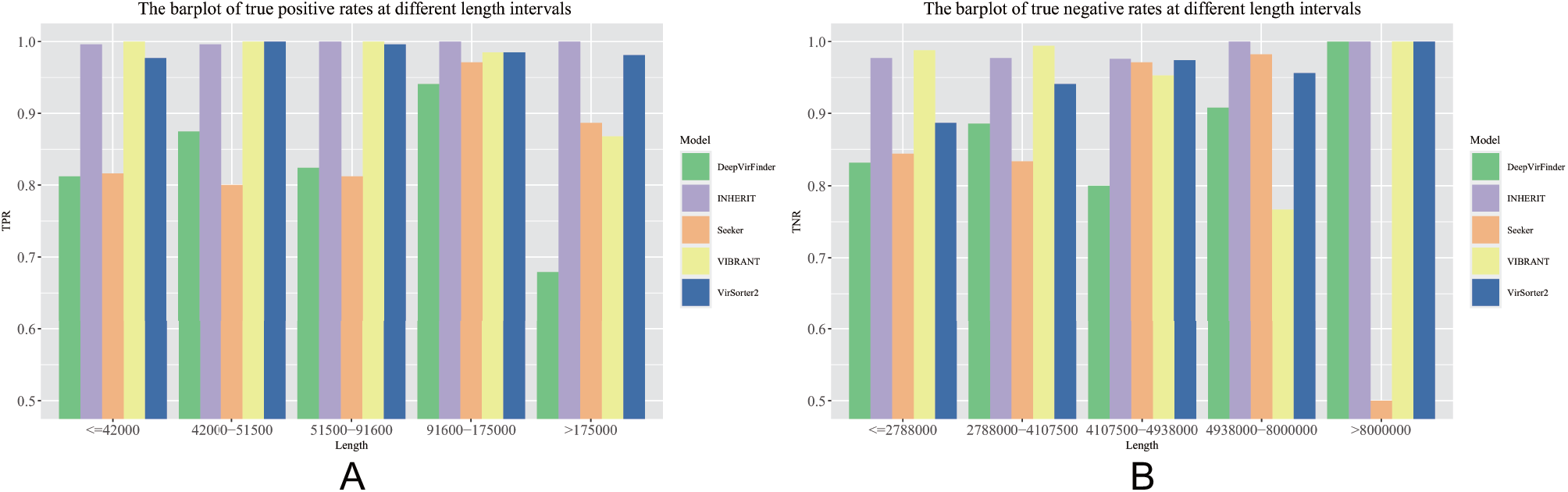
The barplot of the true positive rates and true negative rates at different intervals. (A) shows the true positive rates for each method at different length intervals. (B) shows the true negative rate of each method at different length intervals.

##### INHERIT has appropriate prediction speed

INHERIT is still a model based on a deep learning framework, so it does not take as long a time to predict as database-based approaches. We calculated the average time required for VIBRANT, VirSorter2, Seeker, DeepVirFinder, and INHERIT to predict phage sequences and bacteria sequences in the test set. The results (see Table 2) show that the predictions of VIBRANT and VirSorter2 take much more time than Seeker and INHERIT. From our experiment, VIBRANT and VirSorter2 even cannot predict the whole bacteria test set at once. That indicates that even though database-based methods perform better than alignment-free methods, they consume a long time to predict and are more sensitive to dataset size. However, INHERIT has high performance and predicts the second fastest among VIBRANT, VirSorter, Seeker, and DeepVirFinder. Although INHERIT takes a longer time to predict than Seeker, it takes much less time than DeepVirFinder. DeepVirFinder selects different models for prediction based on the length of the target sequence. In our experiments, we offered the same environment to INHERIT, Seeker, and DeepVirFinder, but DeepVirFinder is still much slower than the other two methods, even if it is based on a convolutional neural network (Lecun and Bottou, 1998; O’Shea and Nash, 2015). We conjecture that it may be because it consumes much time in the process of DNA sequence encoding and model selection. INHERIT uses one model to make predictions for each sequence regardless of the length, showing that INHERIT can give accurate predictions in an appropriate time budget.

**Table 2.** The average time (second) required for VIBRANT, VirSorter2, Seeker, DeepVirFinder, DNABERT, and INHERIT to predict phage sequences and bacteria sequences in the test set. The time of each model implies their average running time (second) on predicting each bacterium and phage. The average length of bacteria samples on the test set is 3,950,500 bp, while the average length of phage samples on the test set is 75,800 bp. Values corresponding to fastest speed are bolded.

| Sequence | VIBRANT | VirSorter2 | Seeker | DeepVirFinder | DNABERT | INHERIT |
| --- | --- | --- | --- | --- | --- | --- |
| Bacteria | 565.7887 | 1256.7568 | **1.6068** | 344.4573 | 53.7031 | 67.3556 |
| Phage | 19.1440 | 29.1797 | **0.1139** | 16.2840 | 2.7851 | 3.0127 |

### 3.2 Ablation study

We used the validation set to do an additional experiment for considering ablation. In this section, we discuss how INHERIT’s pre-training strategy and deep learning structure affect performance.

There are two main differences between INHERIT and the backbone: DNABERT. First, we used a strategy to pre-train two species independently. To use pre-training to provide references and solve imbalanced bacterial and phage datasets, we pre-trained bacteria and phages separately. In addition, we utilized two pre-trained models and have them fine-tuned simultaneously. This model structure has twice the number of parameters as DNABERT. To explore whether pre-training would help deep learning frameworks improve performance, we made an ablation study on the validation set. In training INHERIT, we first trained two pre-trained models on two separate datasets of different sizes and species. Then, the weights of these two pre-trained models were used as initialization while fine-tuning simultaneously on the training set. Therefore, we chose to compare an INHERIT that does not use any pre-trained models and randomly initializes two DNABERTs and fine-tuned them on the training set simultaneously. We denote it as INHERIT (w/o pre-train). Further, since we used two DNABERTs, INHERIT is about twice as large as DNABERT in terms of the number of parameters. Therefore, we also trained a DNABERT to reflect the effect of the number of parameters on the model performance. All models used the same hyperparameters as INHERIT during training.

We tested all models’ performance on the validation set. Here we evaluated all three models on both sequence and segment levels because the validation set is balanced on the segment level while severely imbalanced on the sequence level (43 bacteria and 2,643 phages). The differences in performance reflected by different metrics may be biased when we evaluate them on the sequence level. Therefore, we have attached the confusion matrix of the three models on the level of both sequence and segment in Supplementary Table S3. Based on the results (see Table 3), INHERIT improves on the excellent performance of DNABERT in all metrics and performs better than INHERIT (w/o pre-train) in most of the metrics on sequence level and performs the best on a segment level. Meanwhile, although we trained on segment level, the better performance on the segment level, the more accurate the prediction of the whole genome sequence. The increase in the number of parameters helps INHERIT, allowing it to outperform DNABERT. However, pre-training can give INHERIT a further boost in very high accuracy without increasing the number of parameters. We also made DeLong Test for these three models with each other, and these three curves are statistically significant. Compared to INHERIT (w/o pre-train), INHERIT significantly improves the F1-score from 0.9611 to 0.9943 on the sequence level and from 0.8788 to 0.9542 on the segment level. This reflects that the independent pre-training strategy can help the deep learning framework to distinguish better and make more accurate predictions on both species. Compared with DNABERT, both two differences help INHERIT to make better performance.

**Table 3.** The comparison among DNABERT (without pre-training, i.e., training from scratch), INHERIT (without pre-training), and INHERIT. We use the validation set to evaluate the performance of the three models. Here we evaluate all three models on sequence level and segment because the validation set is balanced on the segment level while severely imbalanced on sequence level. Values corresponding to best performance are bolded.

| Model | Precision | Recall | Accuracy | F1-score | AUROC | AUPRC |
| --- | --- | --- | --- | --- | --- | --- |
| Sequence Level |  |  |  |  |  |  |
| DNABERT | 0.9992 | 0.9224 | 0.9229 | 0.9593 | 0.9751 | 0.9996 |
| INHERIT (w/o pre-train) | **0.9996** | 0.9255 | 0.9263 | 0.9611 | 0.9839 | 0.9997 |
| **INHERIT** | 0.9992 | **0.9894** | **0.9888** | **0.9943** | **0.9971** | **1.0000** |
| Segment Level |  |  |  |  |  |  |
| DNABERT | 0.8767 | 0.8809 | 0.8792 | 0.8788 | 0.9508 | 0.9507 |
| INHERIT (w/o pre-train) | 0.9044 | 0.8951 | 0.9008 | 0.8997 | 0.9530 | 0.9422 |
| **INHERIT** | **0.9436** | **0.9650** | **0.9539** | **0.9542** | **0.9875** | **0.9882** |

Pre-training independently will allow the deep learning framework to give predictions closer to the true values. We plotted boxplots of the scores given by DNABERT, INHERIT (w/o pre-train), and INHERIT for phage and bacteria samples in the validation set on sequence level (see Figure 3). For the vast majority of samples, the score generated by INHERIT is closer to the true value (1 for phages and 0 for bacteria). For example, for phage MH576962, DNABERT gives a score of 0.4707, and INHERIT (w/o pre-train) gives a score of 0.4968. Neither model classify it correctly. However, the score from INHERIT is 0.8182. Compared to the other two models, INHERIT classifies it correctly and significantly improves the score more toward 1. That means that independent pre-training provides great help for deep learning frameworks to give more correct predictions on both species. This point even stands for the samples where none of the three predict correctly or all of the three predict correctly. For example, for phage HQ906662, the DNABERT, INHERIT (w/o pre-train), and INHERIT prediction scores were 0.4209, 0.2908, and 0.4547, respectively; for bacterium NZ_CP018197, the DNABERT, INHERIT (w/o pre-train), and INHERIT prediction scores were 0.2237, 0.1210, and 0.0621, respectively.

**Fig. 3:**
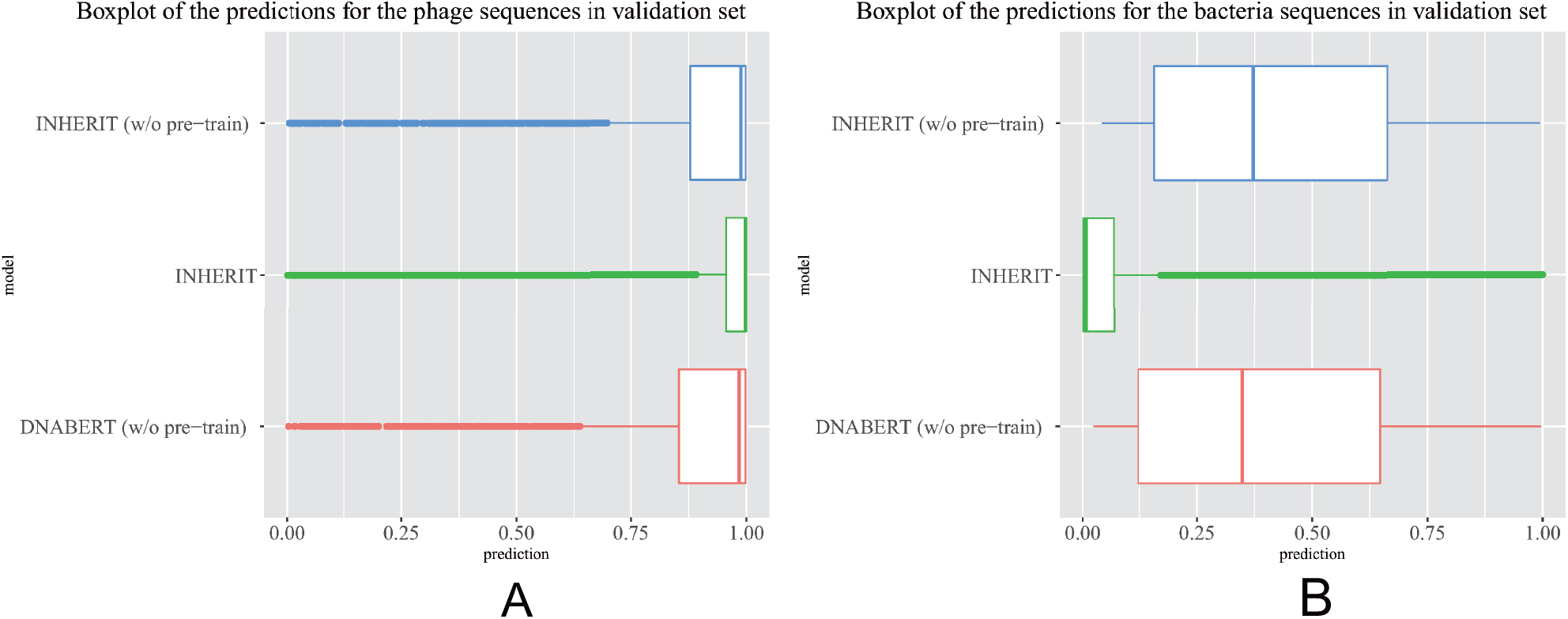
The boxplots of the predictions of DNABERT, INHERIT (w/o pre-train), and INHERIT in the validation set. For phage sequences, the closer the model prediction is to 1, the better. While for bacteria, the closer the model prediction is to 0, the better. (A) is the boxplot of the scores of DNABERT, INHERIT (w/o pre-train), and INHERIT for the phage sequences in the validation set. (B) is the boxplot of the scores of DNABERT, INHERIT (w/o pre-train), and INHERIT for the bacteria sequences in the validation set.

We also tested the prediction speed of DNABERT. From the results (see Table 2), the average prediction speed of DNABERT is 53.7031 seconds for bacteria sequences and 2.7851 seconds for phage sequences in the test set. It is slightly faster than INHERIT. That is because INHERIT has a more complex structure than DNABERT. However, INHERIT can still give predictions faster than other methods.

## 4 Conclusions

In this work, we proposed INHERIT, an integrated method that combines both database-based and alignment-free approaches under a unified deep representation learning framework. It uses two pre-trained models as references and keeps the features of alignment-free methods by the deep learning structure. On a third-party benchmark dataset, we compared the proposed method with VIBRANT, VirSorter2, Seeker, and DeepVirFinder, representing database-based methods and alignment-free methods. We demonstrate that INHERIT can achieve better performance than the four existing methods in all metrics. In particular, INHERIT improves the F1-score from 0.9787 to 0.9932. Meanwhile, we also prove that using an independent pre-training strategy can make deep learning models make better predictions on both species.

## Acknowledgements

The super-computing resource was provided by the Human Genome Center, Institute of Medical Science, The University of Tokyo (https://gc.hgc.jp/en/). Y.Z.’s appreciation also goes to Google who provided GCP based on the partnership between UTokyo and Google.

## Funding

This study was supported by the Ministry of Education, Culture, Sports, Science, and Technology of Japan (Seiya Imoto, 21H03538; Satoshi Uematsu, 21K19495, 22H00477), Japan Society for the Promotion of Science (Y.Z., JSPS KAKENHI Grant Number JP21K12104), the Japan Agency for Medical Research and Development (AMED) (Satoshi Uematsu, 21fk0108619h0001; Kosuke Fujimoto, 21ae0121048h0001), and the Uehara Memorial Foundation (to Seiya Imoto).

### Conflict of Interest

none declared.

The data underlying this article are available in NCBI Nucleotide database at https://www.ncbi.nlm.nih.gov/nuccore/, and the accessions of the sequences used in each dataset can be found in online supplementary materials.

## Notes

### Competing Interest Statement

The authors have declared no competing interest.

### Summary of Updates

Revise the figures and contents minorly.

## References

Auslander, N., Gussow, A. B., Benler, S., Wolf, Y. I., and Koonin, E. V. (2020). Seeker: Alignment-free identification of bacteriophage genomes by deep learning. Nucleic acids research, 48(21), e121–e121.

Bengio, Y., Courville, A., and Vincent, P. (2013). Representation learning: A review and new perspectives. IEEE transactions on pattern analysis and machine intelligence, 35(8), 1798–1828.

Chanishvili, N., Chanishvili, T., Tediashvili, M., and Barrow, P. A. (2001). Phages and their application against drug-resistant bacteria. Journal of Chemical Technology & Biotechnology, 76(7), 689–699.

Chatzou, M., Magis, C., Chang, J.-M., Kemena, C., Bussotti, G., Erb, I., and Notredame, C. (2016). Multiple sequence alignment modeling: methods and applications. Briefings in bioinformatics, 17(6), 1009–1023.

DeLong, E. R., DeLong, D. M., and Clarke-Pearson, D. L. (1988). Comparing the areas under two or more correlated receiver operating characteristic curves: a nonparametric approach. Biometrics, pages 837–845.

Devlin, J., Chang, M.-W., Lee, K., and Toutanova, K. (2018). Bert: Pre-training of deep bidirectional transformers for language understanding. arXiv preprint arXiv:1810.04805.

Dodge, J., Ilharco, G., Schwartz, R., Farhadi, A., Hajishirzi, H., and Smith, N. (2020). Fine-tuning pretrained language models: Weight initializations, data orders, and early stopping. arXiv preprint arXiv:2002.06305.

Eddy, S. R. (1998). Profile hidden markov models. Bioinformatics (Oxford, England), 14(9), 755–763.

Edgar, R. C. and Batzoglou, S. (2006). Multiple sequence alignment. Current opinion in structural biology, 16(3), 368–373.

Edwards, R. A. and Rohwer, F. (2005). Viral metagenomics. Nature Reviews Microbiology, 3(6), 504–510.

Fang, Z., Tan, J., Wu, S., Li, M., Xu, C., Xie, Z., and Zhu, H. (2019). Ppr-meta: a tool for identifying phages and plasmids from metagenomic fragments using deep learning. GigaScience, 8(6), giz066.

Fuhrman, J. A. (1999). Marine viruses and their biogeochemical and ecological effects. Nature, 399(6736), 541–548.

Guo, J., Bolduc, B., Zayed, A. A., Varsani, A., Dominguez-Huerta, G., Delmont, T. O., Pratama, A. A., Gazitúa, M. C., Vik, D., Sullivan, M. B., et al. (2021). Virsorter2: a multi-classifier, expert-guided approach to detect diverse dna and rna viruses. Microbiome, 9(1), 1–13.

Gururangan, S., Marasović, A., Swayamdipta, S., Lo, K., Beltagy, I., Downey, D., and Smith, N. A. (2020). Don’t stop pretraining: adapt language models to domains and tasks. arXiv preprint arXiv:2004.10964.

Ho, S. F. S., Millard, A. D., and van Schaik, W. (2021). Comprehensive benchmarking of tools to identify phages in metagenomic shotgun sequencing data. bioRxiv.

Hyatt, D., Chen, G.-L., LoCascio, P. F., Land, M. L., Larimer, F. W., and Hauser, L. J. (2010). Prodigal: prokaryotic gene recognition and translation initiation site identification. BMC bioinformatics, 11(1), 1–11.

Japkowicz, N. and Stephen, S. (2002). The class imbalance problem: A systematic study. Intelligent data analysis, 6(5), 429–449.

Ji, Y., Zhou, Z., Liu, H., and Davuluri, R. V. (2021). Dnabert: pre-trained bidirectional encoder representations from transformers model for dna-language in genome. Bioinformatics.

Kieft, K., Zhou, Z., and Anantharaman, K. (2020). Vibrant: automated recovery, annotation and curation of microbial viruses, and evaluation of viral community function from genomic sequences. Microbiome, 8(1), 1–23.

Lecun, Y. and Bottou, L. (1998). Gradient-based learning applied to document recognition. Proceedings of the IEEE, 86(11), 2278–2324.

Liu, P., Yuan, W., Fu, J., Jiang, Z., Hayashi, H., and Neubig, G. (2021). Pre-train, prompt, and predict: A systematic survey of prompting methods in natural language processing. arXiv preprint arXiv:2107.13586.

Mao, H. H. (2020). A survey on self-supervised pre-training for sequential transfer learning in neural networks. arXiv preprint arXiv:2007.00800.

Marquet, M., Hölzer, M., Pletz, M. W., Viehweger, A., Makarewicz, O., Ehricht, R., and Brandt, C. (2020). What the phage: A scalable workflow for the identification and analysis of phage sequences. bioRxiv.

Naseem, U., Razzak, I., Khan, S. K., and Prasad, M. (2021). A comprehensive survey on word representation models: From classical to state-of-the-art word representation language models. Transactions on Asian and Low-Resource Language Information Processing, 20(5), 1–35.

O’Shea, K. and Nash, R. (2015). An introduction to convolutional neural networks. arXiv preprint arXiv:1511.08458.

Pedregosa, F., Varoquaux, G., Gramfort, A., Michel, V., Thirion, B., Grisel, O., Blondel, M., Prettenhofer, P., Weiss, R., Dubourg, V., et al. (2011). Scikit-learn: Machine learning in python. the Journal of machine Learning research, 12, 2825– 2830.

Radford, A., Narasimhan, K., Salimans, T., and Sutskever, I. (2018). Improving language understanding by generative pre-training.

Rao, R., Liu, J., Verkuil, R., Meier, J., Canny, J. F., Abbeel, P., Sercu, T., and Rives, A. (2021). Msa transformer. bioRxiv.

Ren, J., Song, K., Deng, C., Ahlgren, N. A., Fuhrman, J. A., Li, Y., Xie, X., Poplin, R., and Sun, F. (2020). Identifying viruses from metagenomic data using deep learning. Quantitative Biology, 8(1).

Reyes, A., Semenkovich, N. P., Whiteson, K., Rohwer, F., and Gordon, J. I. (2012). Going viral: next-generation sequencing applied to phage populations in the human gut. Nature Reviews Microbiology, 10(9), 607–617.

Rodriguez-Valera, F., Martin-Cuadrado, A.-B., Rodriguez-Brito, B., Pasic, L., Thingstad, T. F., Rohwer, F., and Mira, A. (2009). Explaining microbial population genomics through phage predation. Nature Precedings, pages 1–1.

Rohwer, F. and Thurber, R. V. (2009). Viruses manipulate the marine environment. Nature, 459(7244), 207–212.

Thabtah, F., Hammoud, S., Kamalov, F., and Gonsalves, A. (2020). Data imbalance in classification: Experimental evaluation. Information Sciences, 513, 429–441.

Vaswani, A., Shazeer, N., Parmar, N., Uszkoreit, J., Jones, L., Gomez, A. N., Kaiser, Ł., and Polosukhin, I. (2017). Attention is all you need. In Advances in neural information processing systems, pages 5998–6008.

Wolf, T., Chaumond, J., Debut, L., Sanh, V., Delangue, C., Moi, A., Cistac, P., Funtowicz, M., Davison, J., Shleifer, S., et al. (2020). Transformers: State-of-the-art natural language processing. In Proceedings of the 2020 Conference on Empirical Methods in Natural Language Processing: System Demonstrations, pages 38–45.

Zhang, D., Yin, J., Zhu, X., and Zhang, C. (2018). Network representation learning: A survey. IEEE transactions on Big Data, 6(1), 3–28.

