## Supplementary material for "Identification of bacteriophage genome sequences with representation learning": Supplementary Information.docx

Supplementary Methods:

1. Supplement Methods.pdf

Supplement Table:

1. S1: Dataset Information.xlsx: The accessions of the bacterium and phage sequences used on pre-training sets, training sets, validation sets and test sets respectively (Sheet 1: Accessions of the datasets), and their sources on Sheet 2: Data sources.
2. S2: Benchmark Results.xlsx: The predictions and scores for INHERIT, Seeker and VIBRANT on the benchmark test.
3. S3:Confusion Matrix.xlsx: The confusion matrices of DNABERT, INHERIT (w/o pre-train), and INHERIT on validation set on the level of both sequence and segment.
