## Supplementary material for "Identification of bacteriophage genome sequences with representation learning": Supplementary Methods.pdf

Genome Analysis

* To whom correspondence should be addressed.

#### Abstract

**Motivation:** Bacteriophages/Phages are the viruses that infect and replicate within bacteria and archaea. Phages are used to therapeutically provide another potential solution for solving antibiotic resistance, which is one of the threats to global health. To develop phage therapies, the identification of phages from metagenome sequences is the first step. Currently, there are two main methods for identifying phages: database-based (alignment-based) methods and alignment-free methods. Database-based methods typically use a large number of sequences as references; alignment-free methods usually learn the features of the sequences with machine learning and deep learning models.
**Results:** We propose INHERIT which use a deep representation learning model to integrate both database-based and alignment-free methods, combining the strengths of both. Pre-training is used as an alternative way of acquiring knowledge representations from existing databases, while the BERT-style deep learning framework retains the advantage of alignment-free methods. We compare INHERIT with four existing methods on a third-party benchmark dataset. Our experiments show that INHERIT achieves a better performance with the F1-score of 0.9932. In addition, we find that pre-training two species separately helps the non-alignment deep learning model make more accurate predictions.
**Availability:** The codes of INHERIT are now available in: https://github.com/Celestial-Bai/INHERIT.
**Contact:**
**Supplementary information:** Supplementary data are available at *Bioinformatics* online.

### 1 Supplementary Methods

#### 1.1 Related work

**Database-based methods**

This method takes the genome sequence and first predicts its compiled protein using tools such as Prodigal (Hyatt *et al.*, 2010), then compares it with the database by Profile Hidden Markov Models (Eddy, 1998) to determine whether the sequence is a phage. While these methods can generally identify phages with high accuracy, they also have two drawbacks. First, the computational time required to identify phages by these methods is usually long. If a large number of metagenome sequences need to be identified, or if the sequences need to be identified quickly, database-based methods are unsuitable for these situations. At the same time, such methods are primarily limited by the sequences within the reference database, so it is difficult for such methods to identify phages with little sequence similarity to those in the reference databases.

**Alignment-free methods**

This method uses deep learning to extract features directly from DNA sequences to determine whether they are phages. One of the popular approaches is to use Long Short Term Memory (LSTM) (Hochreiter and Schmidhuber, 1997) for training this problem (Liu *et al.*, 2020; Auslander *et al.*, 2020). When the sequences are converted from bases to values, there are usually two ways: one is to convert bases to four values of 1,2,3,4, and the other is to use the one-hot encoding. It uses both strategies to encode and LSTM to classify the sequences. A typical example of this approach is Seeker.

©

| Model name | Type | Input | Output | Commands | Threshold |
|---|---|---|---|---|---|
| VIBRANT | Database-based method | 3-15kb fragments | Assessed over the fragments | python3 VIBRANT_run.py -i sample.fasta | None |
| VirSorter2 | Database-based method | fragments | Assessed over the fragments | virsorter run -w test.out -i test.fa –min-length 1500 -j 4 all | None |
| Seeker | Alignment-free method | 1000 bp-long segments | Asssessed over the sequences | seeker_fasta = SeekerFasta("sample.fasta") predictions = seeker_fasta.phage_or_bacteria() | 0.5 |
| DeepVirFinder | Alignment-free method | No limitation | Assessed over the input | python dvf.py -i ./test/CRC_meta.fa -l 1000 -c 2 | None |
| INHERIT | Integrated method | 500 bp-long segments | Asssessed over the sequences | python3 IHT_predict.py –sequence test_phage.fasta –withpretrain True –model INHERIT.pt –out test_out.txt | 0.5 |

Table 1. Table 4 shows the features of Seeker, VIBRANT, and INHERIT. To make our benchmark test fair, all models used the default commands and hyperparameters to predict our datasets to simulate their performance in most application scenarios. All commands can be found on the GitHub pages of the respective models.

The DNA sequence is changed from a sequence to a matrix with one-hot encoding, so another common idea is to treat this matrix as an image and use Convolutional Neural Network (CNN) (Lecun and Bottou, 1998) to train (Alipanahi *et al.*, 2015). Therefore, researchers proposed DeepVirFinder, which is trained with CNN. It takes not only the original sequences but includes the reverse complemented sequences as input. The other feature is that it chooses the different models to predict the sequence based on its length.

However, the alignment-free methods can only extract some biological features from the training set itself during the training process, but this process of extracting features is limited. Because when we train a deep learning model on a classification task, we usually need an equal or similar amount of positive and negative data (Japkowicz, 2000). However, the number of phages we can obtain from databases is much less than their hosts, bacteria, and the genome sequence lengths of phages usually are much shorter than those of bacteria. Thus, in the past, the number of bacteria selected by these methods tended to be small, which caused information about bacteria to rely too much on these small amounts of bacterial sequence data with a relatively high degree of randomness. Therefore, there is room for improvement in this kind of method.

**DNABERT and our vocabulary-expanded DNABERT**

DNABERT (Ji *et al.*, 2021) is a specific BERT model that modifies the way of tokenizing DNA sequences. BERT (Devlin *et al.*, 2018) stands for Bidirectional Encoder Representations from Transformers and has been widely used in many fields like natural language processing, demonstrating the superiority and power of its structure (Vaswani *et al.*, 2017; Zhou *et al.*, 2021; Dai *et al.*, 2019; Lee *et al.*, 2020; Yang *et al.*, 2019). The success of BERT has also made the pre-train-fine-tune paradigm popular (Gururangan *et al.*, 2020; Radford *et al.*, 2018; Bengio *et al.*, 2013; Lewis *et al.*, 2019). Since BERT can be successful with human language, it is straightforward to think that BERT might also be useful for the language of cells and other biological tissues (i.e., the genome). DNABERT demonstrates the feasibility of this assumption. The authors of DNABERT built pre-trained models with the human genome samples. They achieved state-of-the-art in solving both the human genome and the mammalian genome sequences, demonstrating that the BERT-style frameworks can be used to solve genome-related problems. DNABERT divides the DNA sequence into several tokens by the k-mer encoding so that there will be a finite vocabulary and can be applied to BERT. However, the vanilla DNABERT only considers the four bases without any degenerate bases, which makes it a little challenging to handle sequences containing degenerate bases. Even though there is a proposed solution that Seeker uses a random notation as one of "A", "T", "C" or "G" when dealing with the degenerate bases, the samples themselves will change with the random number if treated this way. Thus, we enlarge the vocabulary of the DNABERT. We unify the degenerate bases as "N" and add them to our vocabulary to ensure fuller information of the sequence read-in. For example, for a sequence "ATCKNTCG", its sequence using 6-mer segmentation is {ATCKNT, TCKNTC, CKNTCG}. We use this kind of vocabulary-expanded DNABERT (noted as "DNABERT" for short in our paper) in all our downstream tasks and experiments.

### 1.2 Hyperparamters and platforms

**Pre-training**: Both pre-trained DNABERT models have the default BERT structure, i.e., 12 hidden layers, 12 attention heads, and 768 embedding size, and since we include the degenerated bases as "N" compared to vanilla DNABERT, the vocabulary of our DNABERT is permutated by five letters (i.e., "A", "T", "C", "G", "N"), with a total of 16,530 words. They are trained on NVIDIA A100 Tensor Core GPUs (40GB memory). Since the sizes of the bacteria pre-training set and the phage pre-training set are different, we choose the different batch sizes and learning rates. We set the batch size to 192 (32 mini-batches per GPU and 6 A100 Tensor Core GPUs) and the learning rate at 5e-4 when pre-training the bacteria dataset while setting the batch size to 20 (10 mini-batches per GPU and 2 A100 Tensor Core GPUs) and learning rate at 1e-3 when pre-training the phage dataset. Both of them use 0.01 weight decay and 10,000 warmup steps. The unsupervised learning task used in the pre-training process of the models is the Masked Language Model. The masked Language Modeling means that some tokens are masked off from the input as they are trained, and then the tokens are predicted by the context. In the BERT model, 15% of the tokens are randomly selected to be masked. 80% of these masked tokens are replaced with [MASK], 10% are replaced with other words, and the remaining 10% keep the original token unchanged. DNABERT still maintains this set of parameters.

**Fine-tuning**: The batch size of the training set is 64, and the validation set is 32, which is the maximum batch size for our device. The learning rate is 1e-5, the maximum learning rate that can be converged, for both pre-trained models, without weight decay and warmup. We also use early stopping based on validation accuracy. This strategy has also been widely used (Chen *et al.*, 2020; Tay *et al.*, 2020; Choromanski *et al.*, 2021; Mostavi *et al.*, 2020). We set the patience to 3, i.e., if the best validation accuracy does not rise in all 3 epochs, the training process will stop. The random seed is set to 6. INHERIT was also fine-tuned under NVIDIA A100 Tensor Core GPUs (40GB memory).

### 1.3 Model setup

The detailed features of the models are shown in Table 4. We use NVIDIA A100 Tensor Core GPUs (40GB memory) to run Seeker, DeepVirFinder, INHERIT, and Intel Xeon Gold 6154 3.0 GHz CPUs (18 cores per CPU) to run VIBRANT and VirSorter2 because they cannot be accelerated with CUDA.

**VIBRANT**: Since VIBRANT is a multi-classifier and its outputs are in segments identified as "organism", "plasmid", and "virus", we need to propose some strategies to make the results consistent.

First, we define the prediction of VIBRANT for each sequence. We choose the prediction with the highest frequency in the segments to which the sequence belongs as the prediction for this sequence. For example, if the predictions of the segments of the target sequence are: "organism", "plasmid", "virus", "virus", we will regard the prediction of VIBRANT for this sequence as a virus. If there are two or more predictions with the highest frequency, we will randomly select one.

Then, since we only identify bacteria and phage sequences, the sequence identified as "organism" and "plasmid" can be considered as "non-phage". In VIBRANT, we equate "non-phage" with the category "Bacteria", scoring with 0, and "virus" with "Phage", scoring with 1. However, these strategies will cause the AUROC and AUPRC of VIBRANT smaller.

We use the default commands to run VIBRANT. However, there were 3 bacteria sequences not given results by VIBRANT, and we did not include them in the calculation of the metrics of VIBRANT.

**VirSorter2**: Since VirSorter2 gives predictions assessed over the segments, we calculate the average of "max_score" of the segment generated by a sequence as the "score" of this sequence. VirSorter2 does not have a default threshold for classifying bacteria and phages. Although Ho et al. give a threshold of 0.93 in their paper, this threshold does not give the optimal results for VirSorter2 in our tests. Therefore, we set the threshold for VirSorter2 to 0.96, i.e., when the score of a sequence exceeds 0.96, it will be identified as a phage, and vice versa, as a bacterium.

We use the default commands to run VirSorter2. However, there were 99 bacteria and 4 phage sequences not given results by VirSorter2, and we did not include them in the calculation of the metrics of VirSorter2.

**Seeker**: We use the default codes and outputs of the Seeker because the outputs of Seeker are consistent with ours.

**DeepVirFinder**: Although DeepVirFinder can predict the sequences directly without segmentation, we cannot run it this way due to memory limitations. Thus, we split the individual sequences into 1000 bp segments (1000 bp is the minimum length for which DeepVirFinder can use the best performance model).

Although the calculation of q-value (Storey, 2003) is given in GitHub by Ren et al. (https://github.com/jessieren/DeepVirFinderexamples)and the classification threshold based on q-value is also given by Ho et al. However, we found that some segments have a p-value of 0 and cannot calculate the q-value according to the GitHub guideline. Moreover, the mean p-value usually does not have meaning in statistics, so we use a mean score of 0.65 as the threshold that can allow DeepVirFinder to achieve the best performance in our tests. When the score of a sequence exceeds 0.65, it will be identified as a phage, and vice versa, as a bacterium.

### References

Alipanahi, B., Delong, A., Weirauch, M. T., and Frey, B. J. (2015). Predicting the sequence specificities of dna-and rna-binding proteins by deep learning. *Nature biotechnology*, **33**(8), 831–838.

Auslander, N., Gussow, A. B., Benler, S., Wolf, Y. I., and Koonin, E. V. (2020). Seeker: Alignment-free identification of bacteriophage genomes by deep learning. *Nucleic acids research*, **48**(21), e121–e121.

Bengio, Y., Courville, A., and Vincent, P. (2013). Representation learning: A review and new perspectives. *IEEE transactions on pattern analysis and machine intelligence*, **35**(8), 1798–1828.

Chen, M., Radford, A., Child, R., Wu, J., Jun, H., Luan, D., and Sutskever, I. (2020). Generative pretraining from pixels. In *International Conference on Machine Learning*, pages 1691–1703. PMLR.

Choromanski, K., Lin, H., Chen, H., and Parker-Holder, J. (2021). Graph kernel attention transformers. *arXiv preprint arXiv:2107.07999*.

Dai, Z., Yang, Z., Yang, Y., Carbonell, J., Le, Q. V., and Salakhutdinov, R. (2019). Transformer-xl: Attentive language models beyond a fixed-length context. *arXiv preprint arXiv:1901.02860*.

Devlin, J., Chang, M.-W., Lee, K., and Toutanova, K. (2018). Bert: Pre-training of deep bidirectional transformers for language understanding. *arXiv preprint arXiv:1810.04805*.

Eddy, S. R. (1998). Profile hidden markov models. *Bioinformatics (Oxford, England)*, **14**(9), 755–763.

Gururangan, S., Marasović, A., Swayamdipta, S., Lo, K., Beltagy, I., Downey, D., and Smith, N. A. (2020). Don't stop pretraining: adapt language models to domains and tasks. *arXiv preprint arXiv:2004.10964*.

Hochreiter, S. and Schmidhuber, J. (1997). Long short-term memory. *Neural Computation*, **9**(8), 1735–1780.

Hyatt, D., Chen, G.-L., LoCascio, P. F., Land, M. L., Larimer, F. W., and Hauser, L. J. (2010). Prodigal: prokaryotic gene recognition and translation initiation site identification. *BMC bioinformatics*, **11**(1), 1–11.

Japkowicz, N. (2000). The class imbalance problem: Significance and strategies.

Ji, Y., Zhou, Z., Liu, H., and Davuluri, R. V. (2021). Dnabert: pre-trained bidirectional encoder representations from transformers model for dna-language in genome. *Bioinformatics*.

Lecun, Y. and Bottou, L. (1998). Gradient-based learning applied to document recognition. *Proceedings of the IEEE*, **86**(11), 2278–2324.

Lee, J., Yoon, W., Kim, S., Kim, D., Kim, S., So, C. H., and Kang, J. (2020). Biobert: a pre-trained biomedical language representation model for biomedical text mining. *Bioinformatics*, **36**(4), 1234–1240.

Lewis, M., Liu, Y., Goyal, N., Ghazvininejad, M., Mohamed, A., Levy, O., Stoyanov, V., and Zettlemoyer, L. (2019). Bart: Denoising sequence-to-sequence pre-training for natural language generation, translation, and comprehension. *arXiv preprint arXiv:1910.13461*.

Liu, F., Miao, Y., Liu, Y., and Hou, T. (2020). Rnn-virseeker: a deep learning method for identification of short viral sequences from metagenomes. *IEEE/ACM Transactions on Computational Biology and Bioinformatics*.

Mostavi, M., Chiu, Y.-C., Huang, Y., and Chen, Y. (2020). Convolutional neural network models for cancer type prediction based on gene expression. *BMC medical genomics*, **13**(5), 1–13.

Radford, A., Narasimhan, K., Salimans, T., and Sutskever, I. (2018). Improving language understanding by generative pre-training.

Storey, J. D. (2003). The positive false discovery rate: a bayesian interpretation and the q-value. *The Annals of Statistics*, **31**(6), 2013–2035.

Tay, Y., Bahri, D., Zheng, C., Brunk, C., Metzler, D., and Tomkins, A. (2020). Reverse engineering configurations of neural text generation models. *arXiv preprint arXiv:2004.06201*.

Vaswani, A., Shazeer, N., Parmar, N., Uszkoreit, J., Jones, L., Gomez, A. N., Kaiser, Ł., and Polosukhin, I. (2017). Attention is all you need. In *Advances in neural information processing systems*, pages 5998–6008.

Yang, Z., Dai, Z., Yang, Y., Carbonell, J., Salakhutdinov, R. R., and Le, Q. V. (2019). Xlnet: Generalized autoregressive pretraining for language understanding. *Advances in neural information processing systems*, **32**.

Zhou, H., Zhang, S., Peng, J., Zhang, S., Li, J., Xiong, H., and Zhang, W. (2021). Informer: Beyond efficient transformer for long sequence time-series forecasting. In *Proceedings of AAAI*.
